# Synergistic elimination of bacteria by phage and the innate immune system

**DOI:** 10.1101/057927

**Authors:** Chung Yin (Joey) Leung, Joshua S. Weitz

## Abstract

Phage therapy has been viewed as a potential treatment for bacterial infections for over a century. Yet, the year 2016 marks the first phase I/II human trial of a phage therapeutic - to treat burn wound patients in Europe. The slow progress in realizing clinical therapeutics is matched by a similar dearth in principled understanding of phage therapy. Theoretical models and *in vitro* experiments find that combining phage and bacteria often leads to coexistence of both phage and bacteria or phage elimination altogether. Both outcomes stand in contrast to the stated goals of phage therapy. A potential resolution to the gap between models, experiments, and therapeutic use of phage is the hypothesis that the combined effect of phage and host immune system can synergistically eliminate bacterial pathogens. Here, we propose a phage therapy model that considers the nonlinear dynamics arising from interactions between bacteria, phage and the host innate immune system. The model builds upon earlier efforts by incorporating a maximum capacity of the immune response and density-dependent immune evasion by bacteria. We analytically identify a synergistic regime in this model in which phage and the innate immune response jointly contribute to the elimination of the target bacteria. Crucially, we find that in this synergistic regime, neither phage alone nor the innate immune system alone can eliminate the bacteria. We confirm these findings using numerical simulations in biologically plausible scenarios. We utilize our numerical simulations to explore the synergistic effect and its significance for guiding the use of phage therapy in clinically relevant applications.

## 1. Introduction

Phage therapy denotes the use of bacteriophage, or viruses that exclusively infect bacteria, as therapeutic agents to treat infections by pathogenic bacteria. Phage therapy was applied in the 1920s and 1930s before the widespread use of antibiotics (Potera, 2013). Early applications of phage therapy were plagued by a number of problems, caused in part by poor understanding of phage biology (Carlton, 1999). These issues range from the lack of proper testing of the host range of phage to inappropriate preparation methods of phage stocks that inactivate the phage. In addition, interest in phage therapy declined after the discovery and mass production of antibiotics. However, the rise of pervasive antibiotic resistance has led to a renewal of interest in using phage to treat bacterial infections (Thiel, 2004).

The first international, single-blind clinical trial of phage therapy, Phagoburn, was launched in 2015 (Clark, 2015). The trial is designed to target 220 burn patients whose wounds are infected by *Escherichia coli* or *Pseudomonas aeruginosa* (Reardon, 2014; Kingwell, 2015). Other clinical trials are currently underway, including a phase I clinical trial by AmpliPhi Biosciences for treating infections of *Staphylococcus aureus* (Kingwell, 2015) and trials targeting respiratory tract infections such as pneumonia (Kingwell, 2015; Fran¸cois *et al*., 2016). Phage-based therapeutics have also been used for *in vitro* biocontrol of pathogenic or nuisance bacteria (Hagens & Loessner, 2007; Abedon, 2009), for example, in the reduction of populations of pathogens in produce (Leverentz *et al*., 2003), ready-to-eat foods (Guenther *et al*., 2009) and meat (Seo *et al*., 2016).

Phage therapy has a number of potential advantages over conventional drug therapy (Loc-Carrillo & Abedon, 2011). Phage can replicate inside bacterial cells and increase in number at the site of infection (Carlton, 1999). This self-amplification is in contrast to the fixed dose in conventional drug therapy (Carlton, 1999; Payne & Jansen, 2000, 2003). A consequence of the self-amplification is that sufficiently high bacteria density is required for the proliferation of phage and the subsequent active control of bacteria (Payne & Jansen, 2000, 2001; Cairns *et al*., 2009). Phage are also more specific than broad spectrum antibiotics. This specificity reduces negative impacts on the normal flora of the human host (Skurnik *et al*., 2007). Some phage also have the ability to break down and clear biofilms which are resistant to antibiotics and recalcitrant to the host immune response (Azeredo & Sutherland, 2008; Rodney, 2009; Timothy & Michael, 2011; Alemayehu *et al*., 2012; Guti´errez *et al*., 2015). Finally, innovations in genetic engineering provide a means to prototype and deploy combinations of phage targeted to individual infections (Campbell, 2003; Hermoso *et al*., 2007).

Despite reinvigorated interest in phage therapy, there remain many questions with respect to how, when, why, and even if phage therapy works. The elimination of a bacterial population by phage is not inevitable, even if phage can eliminate a targeted bacterial population given suitably chosen strains and initial conditions. Indeed, *in vitro* experiments (Chao *et al*., 1977; Levin *et al*., 1977; Lenski, 1984; Dennehy, 2012) and ecological models (Campbell, 1961; Levin *et al*., 1977; Weitz, 2015) of phage-bacteria systems predict broad regimes of coexistence among phage and bacteria populations. In many instances, co-culturing of phage and bacteria together leads to the emergence of resistant host strains and the elimination of the phage population (Dennehy, 2012; Meyer *et al*., 2012). Such outcomes are not desirable from a therapeutic perspective.

Unlike in vitro contexts, the host immune response is a key component of dynamics taking place in a bacterial infection inside animal hosts. Experimental studies have revealed that the host immune response influences the efficacy of phage in clearing pathogenic bacteria in vivo (Tiwari *et al*., 2011; Trigo *et al*., 2013; Pincus *et al*., 2015). The vertebrate immune system is composed of the innate and adaptive immune system. The innate immune response reacts faster than the adaptive system but the latter is more specific and generally more effective. As animal studies of phage therapy have largely focused on acute infections, there is usually insufficient time for the adaptive immune response to fully activate. As such, innate immunity is expected to play a major role in these cases. In particular, neutrophils have been identified as an important player in determining the success of phage therapy for intraperitoneal infection of mice by *P. aeruginosa* (Tiwari *et al*., 2011). In addition, neutrophil dysfunction was found to exacerbate harmful inflammation response induced by excessive phage dosage in a mouse model of Methicillin-resistant *S. aureus* (MRSA) skin infection (Pincus *et al*., 2015). However, for infections by slow-growing bacterium such as *Mycobacterium ulcerans* which has a prolonged clinical course (> 30 days), phage have been shown to stimulate the adaptive immune response by causing increased recruitment of effector T cells to lymph nodes and higher cytokine levels (Trigo *et al*., 2013).

These results highlight the need to understand the role of the immune response in concert with phage therapy. A small number of models have focused on the combined, nonlinear interactions of bacteria, phage, and the immune system (Levin & Bull, 1996, 2004). In such a combined system, phage-bacteria interactions are presumed to operate concurrently with that of the immune response. The immune response eliminates bacteria and the intensity of the immune response is stimulated by the presence of bacteria. Standard models of bacteria-phage-immune system share a few simplifying assumptions (Levin & Bull, 1996, 2004). First, these models do not distinguish between innate and adaptive immunity, which differ in both the time scale of activation and the effectiveness of its antimicrobial action. Second, they assume that the immune response can grow in strength without bounds. Finally, they assume that bacteria cannot evade the immune response. As a result of the latter two assumptions, these models presuppose that bacteria will be eliminated by the immune system when acting alone given sufficient time post-infection. In this framework, the added benefit of phage is to accelerate the time at which the bacteria are eliminated.

Here, we extend earlier models to explore a clinically relevant regime in which bacteria elimination by the immune system is not inevitable, neither in the short- nor long-term. In our model, we demonstrate that the inclusion of immune saturation and bacterial immune evasion are sufficient to give rise to a robust mechanism of effective phage therapy: synergistic elimination of a bacterial pathogen. This synergistic elimination is possible even when neither the innate immune response nor the phage can eliminate the bacterial pathogen, when acting alone. We discuss ways in which this synergistic mechanism can be leveraged in clinical settings.

## 2. Model

**Figure 1:**
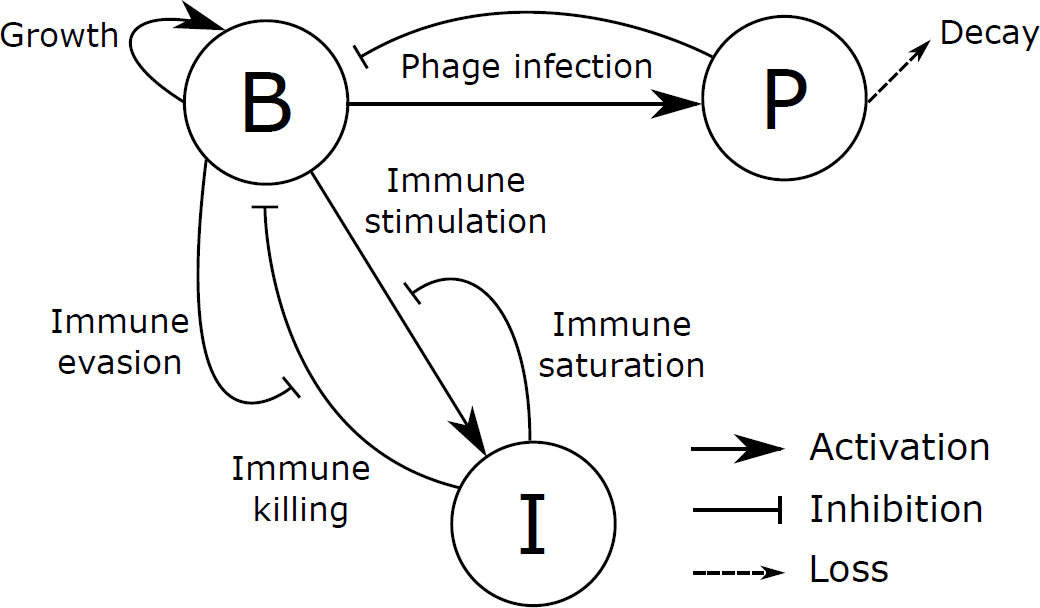
Schematic diagram of a phage therapy model with immune saturation at maximum capacity and immune evasion by bacteria.

We propose a phage therapy model that includes interactions among bacteria B, phage P, and the innate immune response I (see Fig. 1). Here we focus on acute infections and assume only the innate immune system contributes significantly to the killing of bacteria. In this model, the bacteria population reproduces and can be infected and lysed by phage. Phage particles can decay when outside of cells. The presence of bacteria activates the innate immune response which in turn kills the bacteria. This phage therapy model leverages core components proposed in previous models (Levin & Bull, 2004). In so doing, we include two biological features not previously explored.

First, we include a saturating innate immune response. The capacity of the innate immune response is constrained by factors including the finite number of immune cells such as neutrophils that can be produced and tolerated (Summers *et al*., 2010). Experimental studies have shown that host innate immunity has a finite killing capacity that can be saturated (Drusano *et al*., 2010, 2011; Guo *et al*., 2011; Jubrail *et al*., 2016). For example, high bacterial burden can saturate granulocyte killing in a mouse thigh infection model of *Pseudomonas aeruginosa* and *Staphyloccocus aureus* (Drusano *et al*., 2010). This saturation effect allows bacteria to multiply at high densities even under immune killing. The same phenomenon was observed in a murine model of *P. aeruginosa* pneumonia (Drusano *et al*., 2011). A related saturation effect was observed in a murine pneumonia model of *Acinetobacter baumannii* and *P. aeruginosa* (Guo *et al*., 2011) where increase in absolute neutrophil count leads to saturation in bacterial killing rate. In addition, *in vitro* experiments and a murine pneumonia model of *S. aureus* (Jubrail *et al*., 2016) demonstrated that macrophages have a finite capacity for intracellular killing. When this capacity is overwhelmed, bacteria persist intracellularly and are subsequently released through cell lysis. In Fig. 1, this saturation effect is represented as an inhibition of the innate immune activation when the immune response is high, such that there is an upper bound on the magnitude of the host innate immune response.

Second, we include density-dependent immune evasion of bacteria. Experiments have shown that pathogenic bacteria utilize immune evasion strategies such as biofilm formation (Costerton *et al*., 1999) and production of virulence factors (Gellatly & Hancock, 2013) to evade the innate and adaptive immune responses. Coordinated by quorum sensing, bacteria activate these mechanisms at high population density to evade the immune killing action of their host (de Kievit & Iglewski, 2000; Sully *et al*., 2014). As a result, the efficiency of immune killing would decrease with bacterial population density. This is represented as inhibition of the immune killing by bacteria in Fig. 1.

The interactions shown in Fig. 1 are modeled by the following set of differential equations that govern the temporal evolution of bacteria, phage, and the innate immune response:

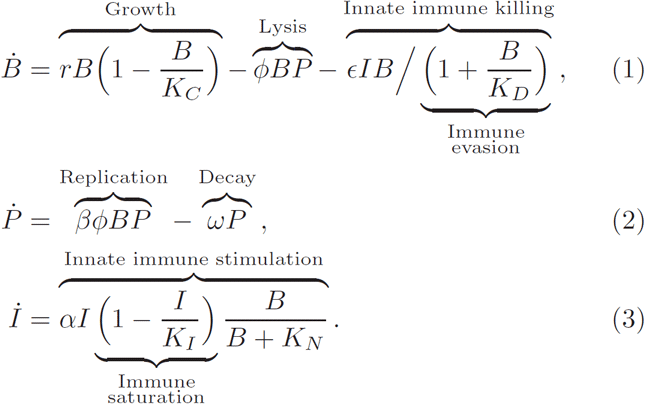

In this model, the maximal bacterial growth rate is r and the carrying capacity is *K_C_*. Phage attach to and infect bacteria at an adsorption rate of *ϕ*, and release new viral particles with a net burst size of β. The phage particles decay or get washed out at a rate of *ω*.

In the absence of immune evasion by bacteria, the immune killing rate in a well-mixed environment is expected to follow the mass action form *ϵIB* where *ϵ* is a killing parameter. To model the effect of immune evasion by bacteria at high densities, we use Michaelis-Menten kinetics where the mass action killing term is scaled by the denominator (1 + *B*/*K_D_*). The saturation results in less efficient innate immune killing at high bacteria density with *K_D_* being the bacteria density at which the host innate immunity is half as effective. This type of kinetics has been used to model immune saturation at high bacterial burden and was found to be in agreement with experimental measurements of *in vivo* bacterial burden (Drusano *et al*., 2010, 2011). The innate immune response is activated by the presence of bacteria with a maximum growth rate of *α*. To model immune saturation, the factor (1 – *I*/*K_I_*) is included to ensure the innate immune response saturates at the immune carrying capacity *K_I_*. Finally, *K_N_* is the bacterial population density at which the innate immune response growth rate is half its maximum. The estimation of model parameters and simulation procedures are described in the Methods section.

## 3. Results

### 3.1. Fixed points of the system and possible infection outcomes

We analyze the fixed points of the system to understand possible outcomes of infection. The fixed points of the system for the non-trivial case of *I* ≠ 0 can be categorized into three different classes (Table 1). The first class has no bacteria but has a finite innate immune response, corresponding to a state in which the innate immune response has eliminated the bacteria. There is also no phage population as phage require bacteria to replicate and persist. In this class, the innate immune response can take on any value less than the maximum, *K_I_*.

The second class of phage-free fixed points contain a nonzero population of bacteria. Here the innate immune response is insufficient to eliminate the bacteria even when it reaches its maximum level, *I*\* = *K_I_*. The equilibrium bacteria density for this class of fixed points depends on the relative magnitude of the bacteria logistic growth rate *G*(*B*) ≡ *r*(1 − *B*/*K_C_*) and the bacteria death rate caused by the maximum innate immune response 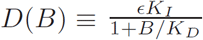. Figure 2 compares the bacterial growth rate with the maximum innate immune killing rate as functions of bacteria density. There is an unstable fixed point 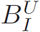 at low bacteria density (main figure) and a stable fixed point 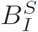 at high bacteria density (inset figure). 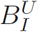 and 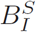 can be obtained by considering *G(B) = D(B)* and solving the resulting quadratic equation, giving

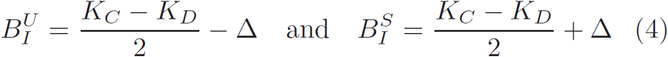

where 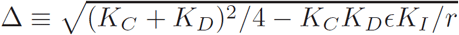.

**Table 1:** The different classes of fixed points of the system and the corresponding infection outcome.

| Class | $B^*$ | $P^*$ | $I^*$ | Outcome |
| --- | --- | --- | --- | --- |
| I | 0 | 0 | $0 < I^* \leq K_I$ | Bacteria elimination |
| II | $0 < B_I < K_C$ | 0 | $K_I$ | Bacteria persistence |
| III | $0 < B_{PI} < K_C$ | $P_{BI} > 0$ | $K_I$ | Coexistence |

When the bacterial concentration is above 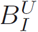, the system will reach the stable fixed point 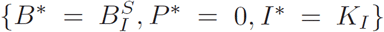 such that the bacteria population persists. This fixed point is reached when there is no phage and the innate immune response alone is insufficient in wiping out the bacteria. On the other hand, when the bacteria density is below 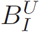, the innate immune killing rate is greater than the bacteria growth rate so the innate immune response can eventually eliminate the bacteria and drive the system to a bacteria-free class I fixed point. Therefore 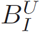 gives the threshold of bacteria density above which the maximum innate immune response fail to eliminate the bacteria. This threshold phenomenon is consistent with experimental observations of the existence of an infective dose required for a bacterial infection to be established (Leggett *et al*., 2012; Smith & Smith, 2016).

Both saturation effects in the model are necessary to produce this threshold phenomenon, i.e., the innate immune intensity must have an upper bound and the efficiency of immune killing must decrease with bacterial burden. Without the first saturation effect (*K_I_* → ∞), the immune response would invariably eliminate the bacteria in the long-term as any fixed point with *B* > 0 and *I* > 0 is impossible (see Eq. (3)). Without the second saturation effect (*K_D_* → ∞), the dynamics would be solely determined by the maximum bacterial growth rate *r* and maximum per capita immune killing rate *ϵK*_*I*_. In such a case, the bacterial population is either always eliminated (r ≤ *ϵK*_*I*_) or never eliminated (r > *ϵK*_*I*_) by the immune response alone, both of which are inconsistent with the experimentally observed threshold phenomenon.

The third class of fixed points exhibits coexistence of bacteria, phage and innate immune response. In this case, the phage persist by infecting and lysing the bacteria population, but the combined effect of phage predation and innate immune response is insufficient to reduce the bacterial population to zero. By Eqs. (1) and (2) the equilibrium bacteria and phage densities are given respectively by

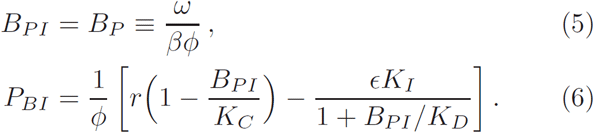

The equilibrium level of bacteria *B_PI_* in the coexistence state depends on the phage parameters and is identical to the equilibrium density *B_P_* when controlled by phage alone. For the coexistence fixed point to be feasible, the phage population density must be positive. A positive phage population occurs when the bacteria growth rate exceeds that of the innate immune killing rate given by *B* = *B*_*P*_. This condition is satisfied when 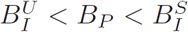.

**Figure 2:**
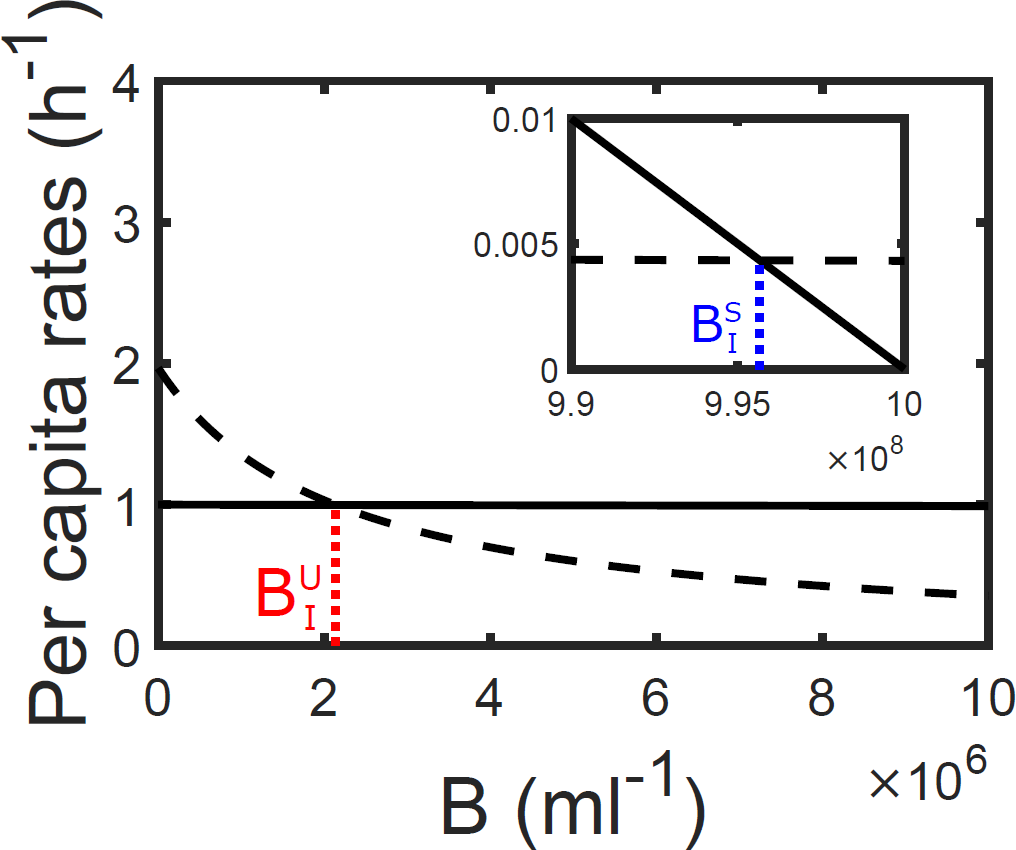
Per capita rates of bacteria growth *G*(*B*) = *r*(1 − *B*/*K_c_*) (solid line) and bacteria death 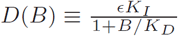 (dashed line) caused by the maximum innate immune response *K_I_* as functions of bacteria density. The inset shows the rates at high bacteria density close to *K_C_*. The red and blue dotted lines mark the positions of 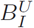 and 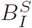 respectively. The model parameters are given in Table 3.

In summary, the bacteria can only be eliminated if the system reaches a class I bacteria-free fixed point. In a class II fixed point, the bacteria persists in the absence of phage. This can happen either when no phage is administered or when the phage administered cannot sustain its population and is eliminated. Finally, the combined killing action of phage and innate immune response is insufficient in eradicating the bacteria in a class III fixed point.

**Figure 3:**
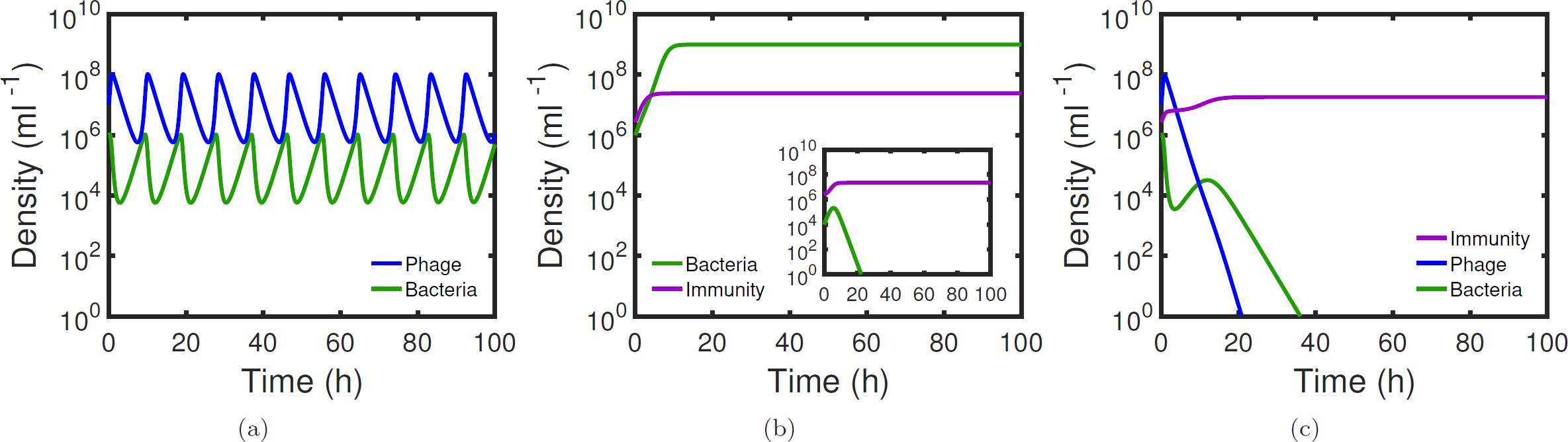
Time series of bacteria density B (green curve), phage density *P* (blue curve) and innate immune response *I* (purple curve) in our model with (a) bacteria and phage only, (b) bacteria and innate immune response only, with the inset showing a lower bacterial inoculum leads to bacterial elimination, and (c) bacteria, phage and innate immune response combined. The model parameters are given in Table 3. The inset in (b) has a lower initial bacterial density of 10^4^ ml^−1^.

### 3.2. Demonstration of a possible synergistic effect between phage and innate immune response

Here we demonstrate that a synergistic effect can be realized in this phage therapy model. In Fig. 3, we compare the dynamics of bacteria, phage, and the innate immune response in three different cases. In the first case (Fig. 3(a)), the bacteria solely interact with phage. The bacteria and phage populations exhibit predator-prey oscillations. These oscillations would eventually be dampened so that populations reach an equilibrium point in the long term (see Eqs. (5) – (6) with *ϵ* = 0) (Weitz, 2015). In this case, the phage alone are unable to eliminate the bacteria. In the second case (Fig. 3(b)), the bacteria interact with the innate immune response in the absence of phage. Initially, there is rapid reproduction of the bacteria and the innate immune response grows as it is stimulated by the presence of bacteria. However, the bacteria persists by reaching a population that is sufficient to evade the maximum innate immune response *K*_*I*_. This is an example of the system converging to a class II fixed point. In this case innate immune response alone is insufficient in eliminating the bacteria. In addition, we show in the inset of Fig. 3(b) that a lower bacterial inoculum can be cleared by the immune response, which is consistent with the existence of an infectious dose required to establish an infection. Finally, in Fig. 3(c) phage and innate immune response act together on the bacterial population initialized above the infectious dose. In this case, oscillations are observed initially for one cycle, but as the innate immune response grows, the phage drive the system to a bacteria-free class I fixed point where the bacteria is eventually eliminated. The phage and innate immune response thus works in synergy to eliminate the bacteria. This synergistic effect is caused by a reduction of the bacterial population by phage below a level of 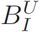. At this level the bacteria are controlled by the innate immune response. With the elimination of the bacterial population, the phage are also eliminated.

### 3.3. Conditions for bacteria elimination through static phage-immune synergy

Next, we simplify the system by applying a quasistatic approximation with the innate immune response treated as a constant. This simplification is justified given that bacteria and phage population are expected to change more rapidly than the innate immune response. We apply this approximation in the case when the innate immune response has reached its maximum *K_I_*. This corresponds to a scenario in which the innate immune response has not controlled a bacterial pathogen. Then phage are added as an additional therapeutic. In this event, the model equations in Eqs. (1) – (3) reduce to

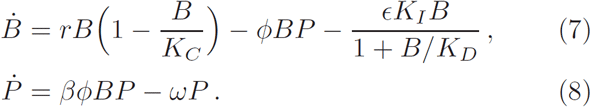

In this approximation, we identify the synergistic effect to mean that phage drive the system originally destined to reach the class II fixed point 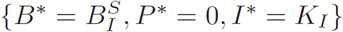 to the class I fixed point 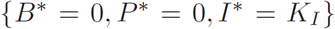 instead. This transition corresponds to the scenario in which phage drive a system from one uncontrolled by the innate immune response to one in which both bacteria and phage are eliminated. This synergistic effect depends on the relative strength of phage and innate immune control of bacteria. The phage-controlled bacteria density is denoted as *B*_*P*_. The bacteria density can reach a maximum of 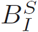 when the innate immune response alone cannot eliminate the bacteria. Synergistic elimination occurs when the phage drive the bacterial population to a level below 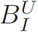 where the maximum innate immune response alone can eliminate the bacteria. This is guaranteed to happen if *B*_*P*_ is below 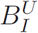, i.e.

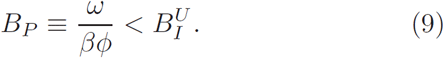

This synergistic effect only relies on the behavior of the system at equilibrium and is independent of the transient dynamics. We therefore denote Eq. (9) as the condition for bacteria elimination through “static synergy”.

In Figs. 4(a) and (b) we show the dynamics of the system with an initial bacteria density *B*_0_ = 10^7^ ml^−1^ and phage dose *P*_0_ = 10^8^ ml^−1^ for a multiplicity of infection (MOI) of 10. The administered phage has an adsorption rate *ϕ* = 10^−8^ ml h^−1^ chosen such that *B_P_* is below 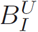. In the absence of innate immune response (*I* = 0), the bacteria and phage populations would oscillate around the fixed point 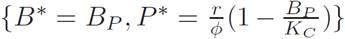 such that part of the trajectory corresponds to states with a bacteria level below 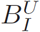. For the set of parameters we consider, 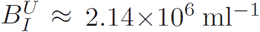 and the condition for static synergy is given by 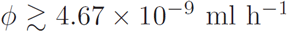. As a result, the bacteria are eliminated given the maximum innate immune response *I* = *K*_*I*_.

It is also important to consider the failure of the synergistic effect. In particular, when 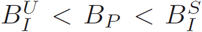, the coexistence steady state given by Eqs. (5) and (6) is feasible. In this event the phage drive the bacteria density to the level *B_P_*. The innate immune response has a marginal effect on the dynamics of the system (Figs. 4(c) and (d)) and no synergistic effect between phage and innate immune response is observed. Finally, when 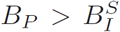 the innate immune response would bring the bacteria density down to 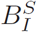 which is insufficient to support the phage population. The phage decay at a faster rate than its replication and eventually die out while the bacteria infection persists.

### 3.4. Conditions for bacteria elimination through dynamic phage-immune synergy

Here, we consider conditions in which the innate immune response alone cannot eliminate the bacteria. We initialize the system in a Class II fixed point. We then consider the effect of adding phage. In Fig. 5 we show simulation results for the steady state bacteria and phage density as functions of the phage decay rate *ω* and adsorption rate *ϕ*. When the phage decay at a slow rate and infects at a high rate which corresponds to a low *B_P_*, it acts synergistically with the innate immune response to eliminate the bacteria. The thresholds 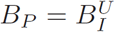 and 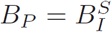 are compared to the simulation results and as expected when 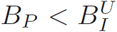 the synergistic effect is always able to exterminate the bacteria. In addition, when 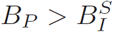 the bacteria persist but phage go extinct. The fixed point analysis predicted a feasible coexistence state of bacteria, phage and innate immune response when 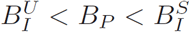, but in the simulation this is only observed for a subset of the region with *B_P_* above an intermediate value. This implies that phage-immune synergy is more robust than expected solely from the feasibility of the coexistence state. This discrepancy can be explained by the stability of the coexistence state. Specifically, bacteria may be eliminated if the coexistence state is unstable. The stability of a dynamical system near a fixed point can be analyzed by considering its Jacobian matrix evaluated at the fixed point (Strogatz, 2014). In Appendix A we show that the condition for stability is:

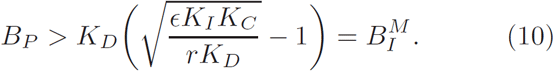

In Fig. 5 we plot the threshold 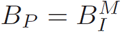 and verify that for 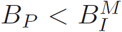 (unstable coexistence state) the bacteria is always driven to extinction.

To show the effect of stability on dynamics, in Fig. 6 we compare the time series of bacteria and phage densities at the three sample points A, B and C from Fig. 5. Case A satisfies 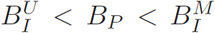 while case B satisfies 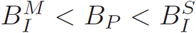. Although coexistence is feasible in both cases, it is unstable in case A and the escalating oscillations caused by any initial deviation from the coexistence state eventually result in extinction of the bacteria. We note that the conditions we derive for instability are sufficient for elimination, and under-estimate the extent to which transient dynamics can enhance the robustness of the synergistic effect. In case B, bacteria persist as the system quickly converges to the coexistence state even from initial conditions far away from it. For case *C*, 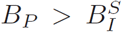 and the coexistence state is infeasible as it has negative phage density. In this case the phage is unable to sustain its population through replication, leading to premature phage extinction.

We summarize the results of the analysis in Table 2. When 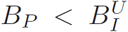 the phage drive the bacterial population below a level that can be controlled by the innate immune response which leads to bacteria elimination by static phage-immune synergy. When 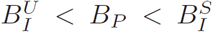, the phage can coexist with the bacteria in the presence of innate immune response, but when 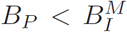 the coexistence is unstable and the oscillations can still drive the bacteria to extinction via a dynamic synergistic effect. Finally, when 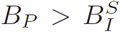 the phage become extinct rapidly as the bacteria persist at a level which cannot support phage growth. We conclude that the synergistic effect is enhanced when taking into account the stability of the coupled, nonlinear dynamical system. In Appendix B we show that the general results of the analysis and the enhanced synergy domain are robust to model formulations. We do so by extending the model to include an explicitly infected class of bacteria. In this scenario, we still find the same qualitative results as in the model where infection is modeled implicitly (see Fig. B.8). In addition, we investigate the effect of initial conditions on the success of phage therapy in Appendix C. We show that phage-immune synergy is robust to initial bacterial inoculum and phage dose.

**Figure 4:**
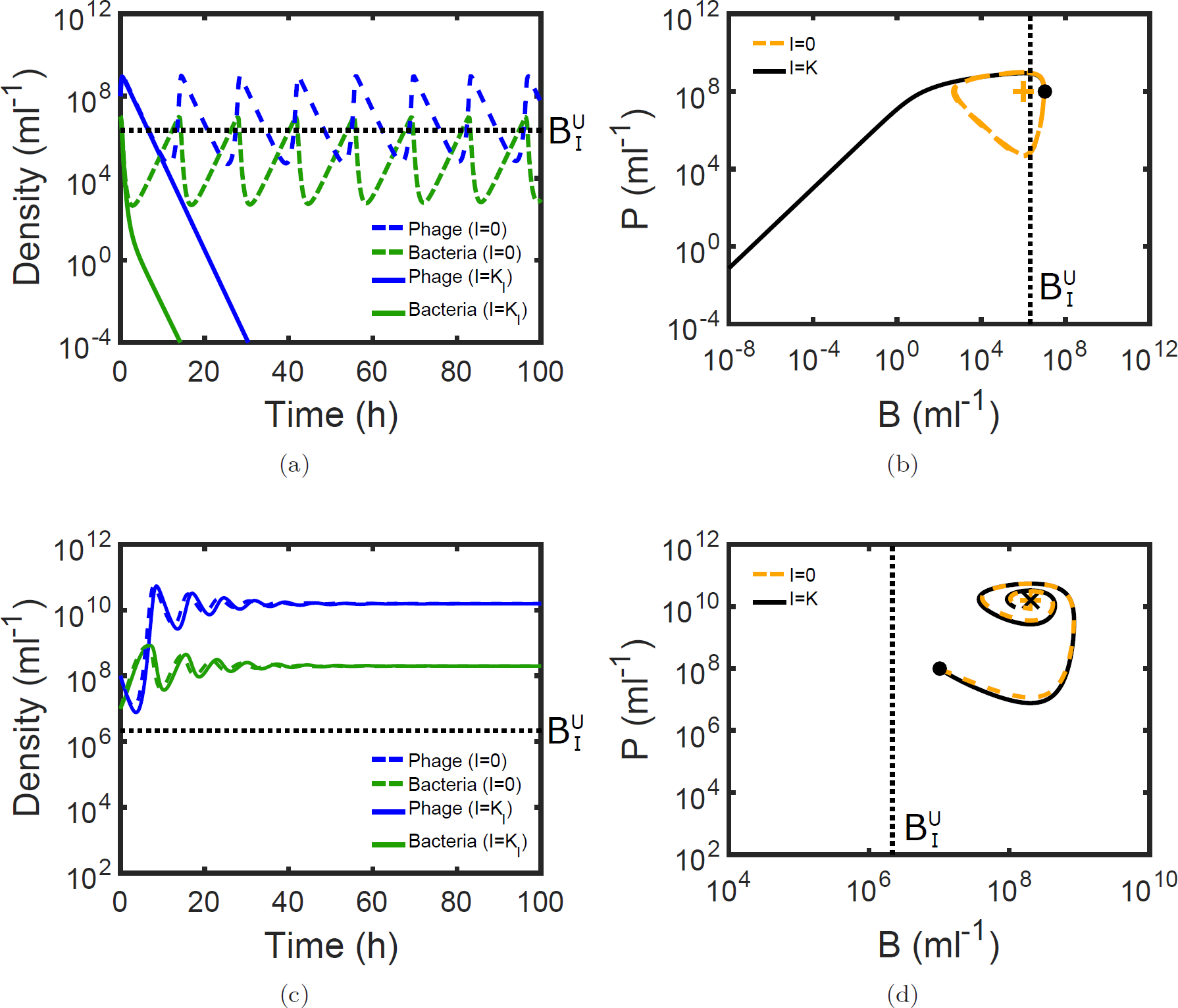
Comparison of time series and phase portraits with and without innate immune response for two different adsorption rates. (a) Time series of bacteria (green curves) and phage densities (blue curves) with adsorption rate *ϕ* = 10^−8^ ml h^−1^. Solid lines are for the case of *I* = *K_I_* while dashed lines are for the case without innate immune response (I = 0). The horizontal dotted line mark the position of 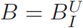, the bacteria density below which the maximum immune response alone can eliminate the bacteria. (b) shows the phase portrait for the time series in (a). Black solid line is the phase trajectory when *I* = *K_I_* while orange dashed line is the trajectory when *I* = 0. The vertical dotted line mark the position of 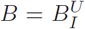 as in (a). The initial conditions (*B*_0_ = 10^7^ ml^−1^, *P*_0_ = 10^8^ ml^−1^) of the trajectory is marked by a filled circle and the fixed point 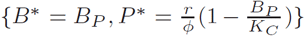 without innate immunity (*I* = 0) is denoted by a plus sign. (c) and (d) are the same as (a) and (b) but with a lower adsorption rate *ϕ* = 5 × 10^−11^ ml h^−1^. The coexistence fixed point {*B*\* = *B_PI_, P*\* = *P_BI_, I*\* = *K_I_*} is marked by a cross in (d).

**Figure 5:**
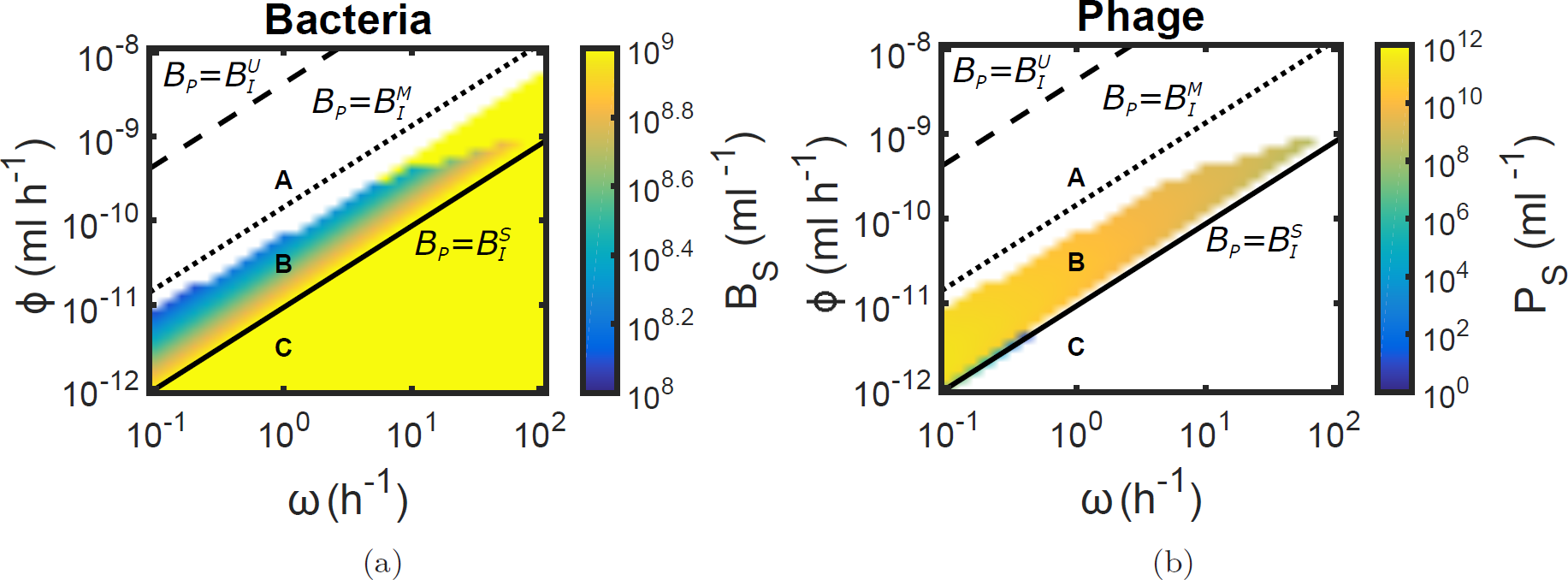
Heat map showing the dependence of (a) log steady state bacteria density *B_S_* and (b) log steady state phage density *P_S_* on the phage decay rate *ω* and adsorption rate *ϕ*. The white region represents the parameter space with (a) elimination of bacteria and (b) phage extinction. The solid line is 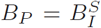, the dashed line is 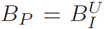 and the dotted line is 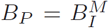. The points labeled by A, B and C are sample points of the different regimes with time series shown in Figs. 6(a), (b) and (c) respectively. The initial conditions are given by *B*_0_ = 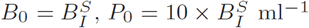 and *I*_0_ = *K*_*I*_.

**Figure 6:**
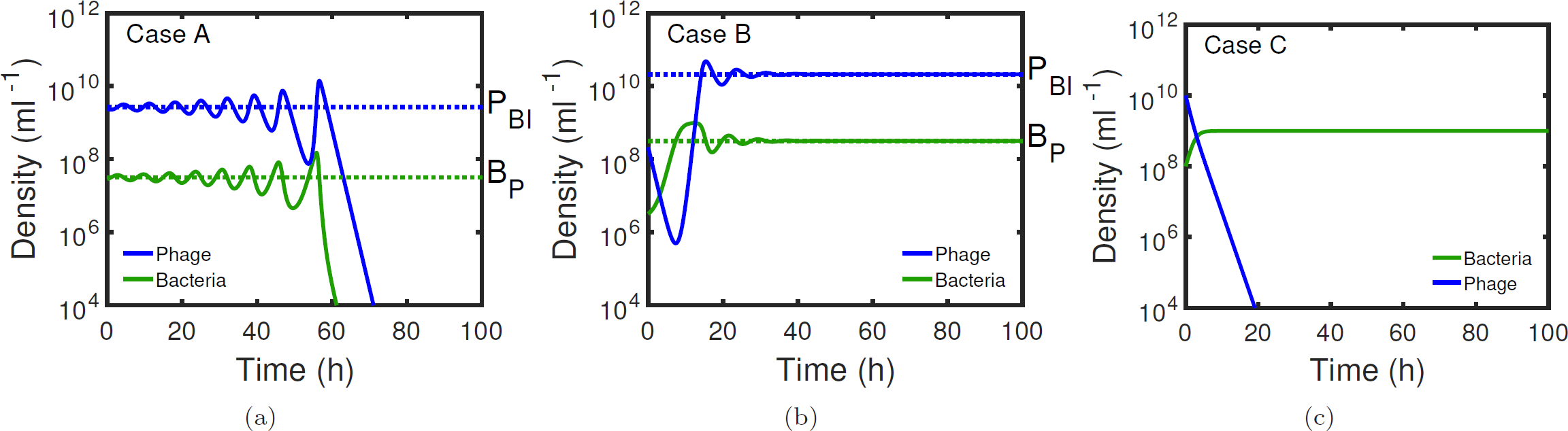
Time series of bacteria (green curve) and phage densities (blue curve) at the three sample points A, B and C in Fig. 5: (a) *ϕ* = 10^−9.5^ « 3.16 × 10^−10^ ml h^−1^ 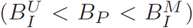, (b) *ϕ* = 3.16 × 10^−11^ ml h^−1^ 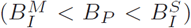 and (c) *ϕ* = 3.16 × 10^−12^ ml h^−1^ 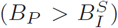. The phage decay rate *ω* =1 h^−1^ for all three cases. The initial conditions for (a) and (b) are chosen by perturbing from the coexistence state, i.e. (*B*_0_ = 0.9 × *B_P_*(*ϕ*), *P*_0_ = 0.9 × *P_BI_*(*ϕ*)) and (*B*_0_ = 0.01 × *B_P_*(*ϕ*), *P*_*0*_ = 0.01 × *P_BI_*(*ϕ*)) for (a) and (b) respectively. For (c) where the coexistence state is infeasible, the initial conditions are *B*_0_ = 10^8^ ml^−1^ and *P*_0_ = 10^10^ ml^−1^.

**Table 2:**
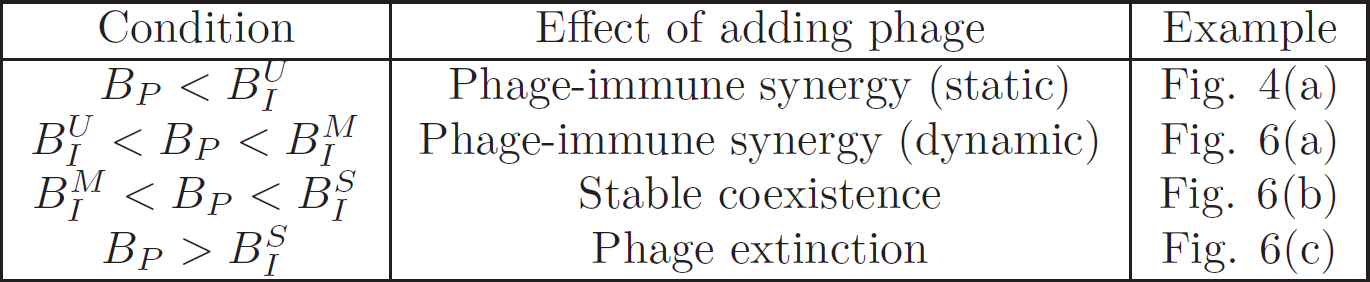
Different regimes of the effect of phage therapy, the corresponding conditions and examples of time series in each regime.

| Condition | Effect of adding phage | Example |
| --- | --- | --- |
| $B_P < B_I^U$ | Phage-immune synergy (static) | Fig. 4(a) |
| $B_I^U < B_P < B_I^M$ | Phage-immune synergy (dynamic) | Fig. 6(a) |
| $B_I^M < B_P < B_I^S$ | Stable coexistence | Fig. 6(b) |
| $B_P > B_I^S$ | Phage extinction | Fig. 6(c) |

## 4. Discussion

The science of phage therapy remains controversial despite renewed interest in translational and clinical applications. In particular, the interactions between bacteria, phage and the host immune response has yet to be fully characterized. Building on existing model of phage-bacteria-immune system (Levin & Bull, 2004), we developed a model of phage therapy that takes into account the finite capacity of host innate immune response and immune evasion of bacteria at high population densities. In this model, we identified a potential mechanism of phage therapy that relies on a synergistic effect between phage and the host innate immune response instead of through direct elimination of bacteria by phage. We show that the synergistic effect is robust to model formulation, bacterial inoculum, and phage dosage. Our results are qualitatively consistent with the experimental observation that phage treatment is effective in protecting normal mice from bacterial infection but ineffective for neutropenic mice with a suppressed innate immune response (Tiwari *et al*., 2011).

The synergistic effect of phage therapy can be understood intuitively via the schematic in Fig. 7. In the initial phase of the infection, a small population of bacteria invade and reproduce within the human host. The bacterial population can become large enough to utilize immune evasion strategies such as biofilm formation if its growth is sufficiently fast compared to the activation of the innate immune response, for example, via production and recruitment of innate effector cells such as neutrophils. In such a case, the biofilm protects the bacteria from phagocytosis by the innate effector cells (Costerton *et al*., 1999), thus the innate immune response cannot eliminate the bacteria.

Addition of phage can reduce densities, e.g., by breaking down density-dependent defenses like biofilms (Azeredo & Sutherland, 2008; Rodney, 2009; Timothy & Michael, 2011; Alemayehu *et al*., 2012; Gutiérrez *et al*., 2015). The resulting reduced bacterial population exposes the bacteria to the action of the innate immune response which eventually eliminates the harmful bacteria. The phage population is eventually eliminated as its target hosts are eliminated. The model formulation is relevant to density-dependent bacterial inhibition of immune clearance, whether via biofilm formation or other means.

We have derived the conditions that predict the outcome of phage therapy in this model. The conditions for effective therapy depend on the relative strength of the phage and immune control of bacteria. Phage with a sufficiently high effectiveness act synergistically with the innate immune response in either a static or dynamic manner. In a static phage-immune synergy, phage lower the bacterial population to a point where the innate immune response can then eliminate it. In a dynamic phage-immune synergy, the introduction of phage destabilizes the system and the subsequent oscillations drive the bacteria densities sufficiently low to be eliminated by the immune response. Phage with intermediate effectiveness can stably coexist with the innate immune response without clearing the bacteria infection. Phage with low effectiveness will become extinct prematurely. The analytical conditions for phage therapy success also provide insights on the type of phage that would be effective. Our results indicate that phage with a low value of *B_P_* = *ω*/(*βϕ*) would be more effective as a therapeutic agent, i.e., a low decay rate, high adsorption rate and large burst size. Importantly, the condition suggests that multiple factors have to be considered when evaluating the suitability of a phage strain for therapeutic use. For example, a phage strain with relatively low adsorption rate may still be effective if it has a long biological half-life *in vivo*.

**Figure 7:**
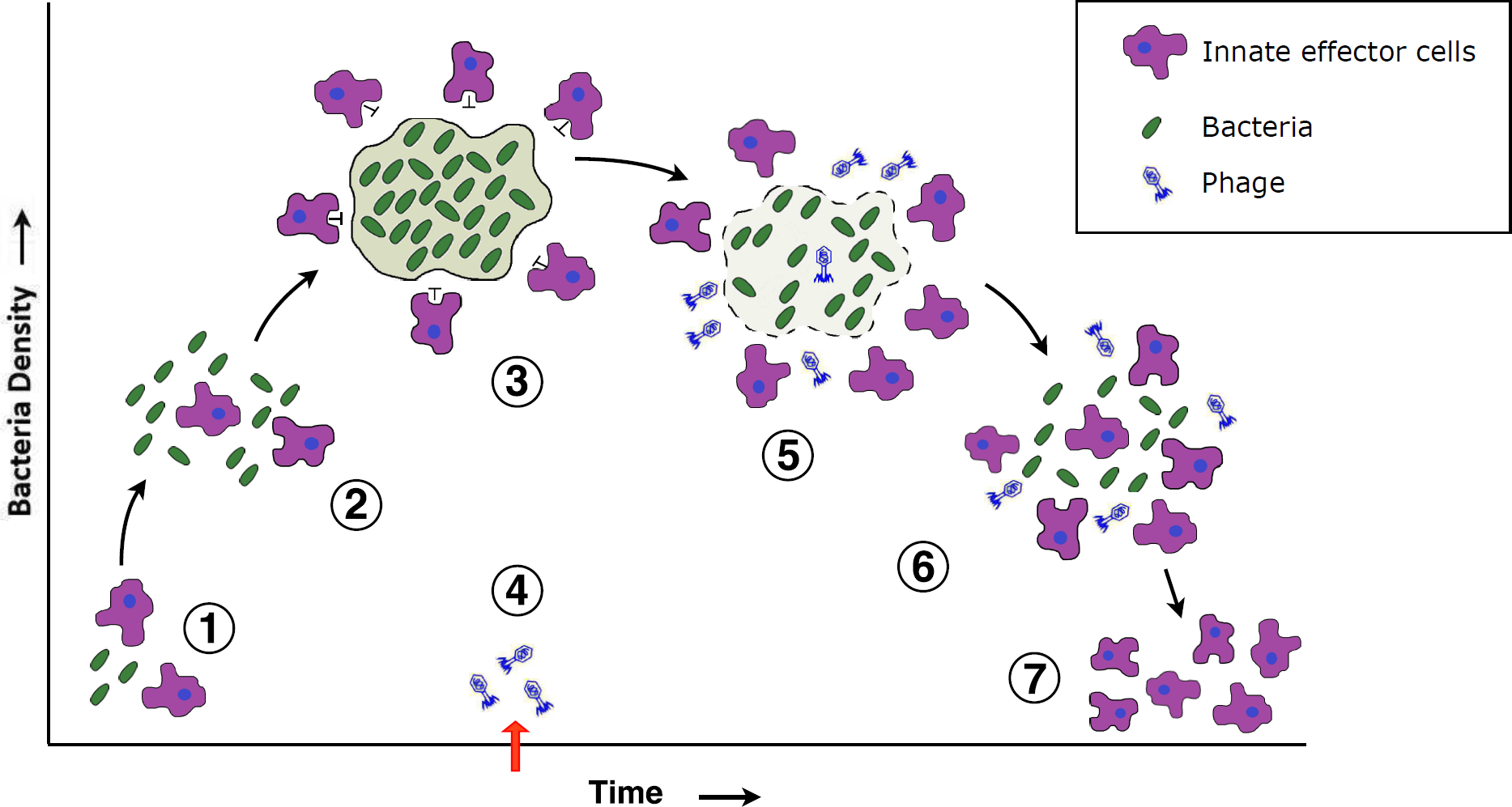
Schematic of the mechanism of phage-immune synergy. Various stages of the infection and therapy: 1. Invasion of human host by bacteria, 2. Reproduction of bacteria, 3. Bacteria form a biofilm while innate immune response gets activated via production and recruitment of innate effector cells, 4. Administration of phage, 5. phage break down biofilm and reduce bacterial population, 6. Innate effector cells gain access to bacteria, and 7. Elimination of bacteria by innate effector cells and phage. Phage are eventually eliminated due to the absence of bacteria.

The proposed model enables analytical solutions for the conditions under which the synergistic effect can eliminate the bacteria. These conditions facilitate the identification of general principles of the synergism. However, the simplification also results in certain limitations which will have to be addressed in future developments of the model. For example, our simulation results indicate that the time scale of the immune response is important in determining the outcome of infection and phage administration. In our model we have focused on the innate immune system, which has a faster time scale than the adaptive immune system but is also generally less effective in comparison. As a result, the adaptive immune response may be more critical in determining the efficacy of phage therapy in chronic infections. Future modeling efforts should explore the existence of multiple time scales and immune response strengths on the outcome of phage therapy.

Another complication that may arise in the application of phage therapy is the development of phage resistance in bacteria. Bacteria employs a wide range of strategies to evade phage infection ranging from modification of surface receptors to degradation of exogenous genetic elements by mechanisms such as the CRISPR-Cas system (Labrie *et al*., 2010). Phage can also evolve counter-resistance and restore infectivity against resistant strains of bacteria, resulting in coevolution between the phage and bacteria (Buckling & Rainey, 2002; Weitz *et al*., 2005; Hyman & Abedon, 2010). Experiments have revealed that the co-evolutionary dynamics can be categorized into at least two contrasting types: arms race dynamics where the bacteria becomes increasingly resistant while the phage become more infective, and fluctuating selection dynamics where there is no directional change in the bacteria resistance range and phage host range (Hall *et al*., 2011; Betts *et al*., 2014). Development of phage resistance in bacteria also alters the way they interact with the immune system and in many cases will lead to reduced virulence (Filippov *et al*., 2011; Seed *et al*., 2014). The effects of the coevolutionary dynamics between bacteria, phage and immune response on the efficacy of phage therapy still remains unclear. Generalizing phage therapy models to include multiple evolved strains of bacteria and phage will be essential.

Finally, it has been recently suggested that phage may be removed or degraded by both the innate and adaptive arms of the host immune system (Hodyra-Stefaniak *et al*., 2015). Such removal may affect the clinical outcome of phage therapy. This effect is assumed to be constant in our model and incorporated into the decay rate *ω*. However, the immune suppression of phage may increase as the bacteria activate the innate immune response. This can be modeled as a phage decay rate *ω*(*I*) that depends on the immune response. In such a case the outcome of phage administration is likely to depend on the relative strength between this direct immune inhibition of phage and the phage-immune synergistic effect.

How can we translate model findings into a therapeutic context? One finding of our model is that phage that are only partially effective in killing bacteria *in vitro* may still be effective in vivo due to the synergistic effect with the host innate immune response. Current screening of phage strains for phage therapy relies on *in vitro* killing (Gill & Hyman, 2010; Khan Mirzaei & Nilsson, 2015). As a consequence, phage strains of therapeutic value may be overlooked. The synergistic effect observed in our model highlights the need to measure the interactions between bacteria, phage and components of the immune system both *in vitro* and *in vivo*. Such experiments complement prior studies involving phage and bacteria (Levin *et al*., 1977; Buckling & Rainey, 2002; Brockhurst & Koskella, 2013) and between bacterial pathogens and innate effector cells such as neutrophils (Li *et al*., 2002) and macrophages (Nau *et al*., 2002; Ensminger *et al*., 2012). Information collected from such tripartite experiments would be important to understand the effects of phage on microbes in animal hosts and within clinical studies.

## 5. Methods

### 5.1. Model simulation

We simulate the model in Eqs. (1) – (3) and obtain the time series of the bacteria density, phage density and innate immune response intensity. The numerical integration is carried out using ode45 in MATLAB. A threshold for the population densities is implemented such that when the bacteria or phage densities fall below a set value *p*_*th*_ = 1 ml^−1^, it is assumed to be extinct and the population density is set to 0. The steady state bacteria level B_S_ is estimated by running the simulation for a sufficiently long time (*T* = 10000 h) before averaging over a duration of *T*_*av*_ = 1000 h. The code for our simulations is available online^1^.

### 5.2. Parameter estimation

In Table 3 we show estimated values of the parameters in the model which are used in the simulations unless otherwise specified. The life history traits of the bacteria and phage are obtained from well-established empirical values for common bacterial pathogens and their phage in the literature (Guo *et al*., 2011; Capparelli *et al*., 2010; De Paepe & Taddei, 2006; Shao & Wang, 2008). The decay rate ω of phage is estimated from measured clearance of phage from the blood of mice (Hodyra-Stefaniak *et al*., 2015) and is higher than typical decay rates of phage *in vitro* (De Paepe & Taddei, 2006).

The maximum growth rate of the innate immune response *α*, the initial (baseline) innate immune intensity *I*_0_, and the maximum innate immune response are estimated from measurements of neutrophil recruitment in LPS-induced acute lung injury (Reutershan *et al*., 2005). Since LPS is highly immunogenic, we assume that it saturates the antigen dose response of the immune activation (*B* ≫ *K*_*N*_). Under this limit, Eq. (3) yields

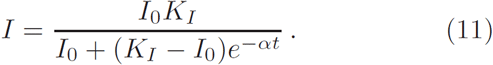

We normalize the measured interstitial neutrophil count (Reutershan *et al*., 2005) of the whole lung by the estimated lung tissue volume of 0.135 ml to obtain the time series of neutrophil concentration. We then perform a nonlinear fitting of Eq. (11) to the time series data using the damped least-squares method computed with the Grace graphing package and obtain the parameter values.

The killing rate parameter of the innate immune response e is estimated by setting it such that *ϵK*_*I*_ gives the fitted maximum killing rate of neutrophils in a murine model of pneumonia caused by *P. aeruginosa* (Drusano *et al*., 2011). We utilize the same dataset to obtain the bacterial concentration K_*D*_ at which innate immune response is half saturated (Drusano *et al*., 2011). The bacteria density *K_N_* at which innate immune growth rate is half its maximum is estimated from the effect of bacterial burden on macrophage recruitment rate in *Mycobacterium tuberculosis* infection (Wigginton & Kirschner, 2001; Sadek *et al*., 1998). Initial bacterial load is set by typical values of bacterial inoculum for *P. aeruginosa* and *S. aureus* (Drusano *et al*., 2010, 2011). The initial phage dose is selected to give a multiplicity of infection (MOI) of 10 which was determined by experiments to provide optimal effectiveness for respiratory infections by *P. aeruginosa* (Debarbieux *et al*., 2010) and *Burkholderia cepacia* complex (Semler *et al*., 2014).

**Table 3:** Estimated values of parameters in the model

| Symbol and parameter | Value | Estimated from |
| --- | --- | --- |
| $r$, growth rate of bacteria at low density | $1 \text{ h}^{-1}$ | *A. baumannii* and *P. aeruginosa* (Guo *et al.*, 2011) |
| $K_C$, carrying capacity of bacteria | $10^9 \text{ ml}^{-1}$ | Bacterial load of *P. aeruginosa* in lung tissue of immunosuppressed mice (Guo *et al.*, 2011) |
| $\beta$, burst size of phage | 100 | Phi1 phage in *Salmonella* (Capparelli *et al.*, 2010) and within the range for coliphage (De Paepe & Taddei, 2006) |
| $\phi$, adsorption rate of phage | $5 \times 10^{-8} \text{ ml h}^{-1}$ | Within the range for Lambda phage (Shao & Wang, 2008) and other coliphage (De Paepe & Taddei, 2006) |
| $\omega$, decay rate of phage | $1 \text{ h}^{-1}$ | Clearance of phage from blood of mice (Hodyra-Stefaniak *et al.*, 2015) |
| $\alpha$, maximum growth rate of innate immune response | $0.97 \text{ h}^{-1}$ | Fitting of interstitial neutrophil recruitment data in LPS-induced acute lung injury (Reutershan *et al.*, 2005) |
| $I_0$, initial innate immune intensity | $2.7 \times 10^6 \text{ (ml}^{-1}\text{)}$ | Fitting of neutrophil recruitment data in acute lung injury (Reutershan *et al.*, 2005) |
| $K_I$, maximum capacity of innate immune response | $2.4 \times 10^7 \text{ ml}^{-1}$ | Fitting of neutrophil recruitment data in acute lung injury (Reutershan *et al.*, 2005) |
| $\epsilon$, killing rate parameter of innate immune response | $8.2 \times 10^{-8} \text{ ml h}^{-1}$ | Set such that $\epsilon K_I$ gives $1.97 \text{ h}^{-1}$, the fitted maximum killing rate of neutrophils in a murine model of pneumonia (Drusano *et al.*, 2011) |
| $K_D$, bacterial concentration at which innate immune response is half saturated | $2.2 \times 10^6 \text{ ml}^{-1}$ | Fitted parameter in a murine model of pneumonia by *P. aeruginosa* (Drusano *et al.*, 2011) |
| $K_N$, bacterial concentration when innate immune response growth rate is half its maximum | $10^5 \text{ ml}^{-1}$ | Effect of bacterial burden on macrophage recruitment rate in *Mycobacterium tuberculosis* infection (Wigginton & Kirschner, 2001; Sadek *et al.*, 1998) |
| $B_0$, initial bacteria density | $10^6 \text{ ml}^{-1}$ | Typical bacterial inoculum used for *in vivo* growth experiments of *P. aeruginosa* and *S. aureus* (Drusano *et al.*, 2010, 2011) |
| $P_0$, initial phage dose | $10^7 \text{ ml}^{-1}$ | Set to give a multiplicity of infection (MOI) of 10 required to ensure effective phage therapy (Debarbieux *et al.*, 2010; Semler *et al.*, 2014) |

## Appendix A. Derivation of the stability condition for the coexistence state

Here we derive the condition for the coexistence state {*B*\* = *B*_*P*_, *P*\* = *P*_*BI*_} to be stable. The type and stability of a fixed point can be characterized by the determinant *det*(**J**) and trace *tr*(**J**) of the Jacobian matrix **J** evaluated at the fixed point (Strogatz, 2014). In particular, the stability of the system is determined by the signs of *det*(**J**) and *tr*(**J**).

Rewriting Eqs. (7) and (8) into 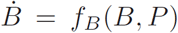 and 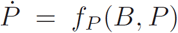 respectively, the Jacobian matrix of the system is defined as

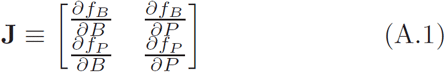

We compute the Jacobian matrix by calculating the partial derivatives:

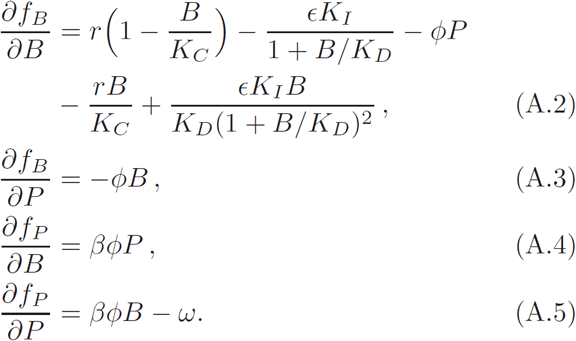

The Jacobian matrix of the system can therefore be written as

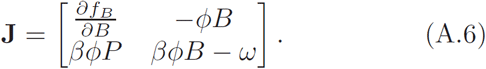

At the coexistence state, the first line on the right hand side of Eq. (A.2) vanishes as a result of Eq. (6). Together with *βϕB*_*P*_ − *ω* = 0, it yields the Jacobian matrix at co-existence

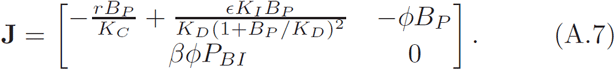

Equation (A.7) gives the following determinant and trace:

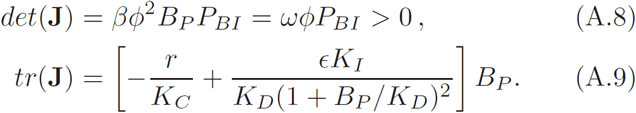

Since the determinant is always positive, the condition for stability is given by *tr***J** < 0 (Strogatz, 2014). By Eq. (A.9) this is satisfied when

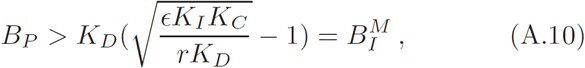

which is the criterion in Eq. (10).

## Appendix B. Bacteria-phage-immune model with an infected bacterial class

In this appendix we show that the general results of our model is robust to variations in model formalism. In our model, the time between adsorption of a phage particle to lysis of the bacterial cell, or the latent period, is assumed to be sufficiently small such that the replication and release of new phage particles can be treated as instantaneous. Here we extend the model to account for the latent period by introducing an infected bacterial class *F* explicitly in which individuals are lysed at a rate *η* (Weitz, 2015). The generalized model is given by

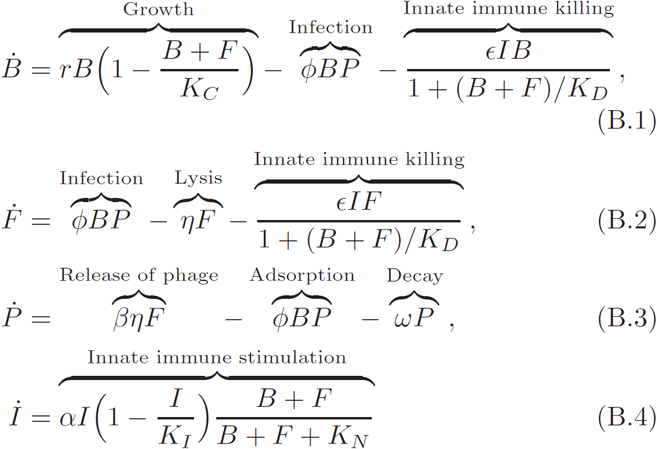

where *F* is the population density of the bacteria infected by phage, *η* is the rate at which the infected bacteria lyse, and the rest of the parameters are defined in the original model in Eqs. (1) – (3).

We simulate this model by fixing *I* = *K*_*I*_ as in Fig. 5 and show the steady state populations of uninfected bacteria, infected bacteria and phage in Fig. B.8. The results show all the same qualitative features as in Fig. 5 including regimes of phage-immune synergism, phage-bacteria-immune coexistence and phage extinction. This indicates that the results of our analysis is robust to the model formulation. The results differ from the model without an infected class only by a scaling of the thresholds 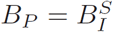, 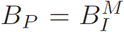 and 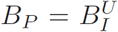. The scaling is caused by a reduction of the effective burst size that results from infected bacteria being killed by the innate immune response before the phage particles are released. As a result, for the phage infection to successfully produce viral progeny, the lysis must be completed before the bacterial cell is killed by the innate immune response. Assuming the innate immune response is at its maximum *K*_*I*_, this gives the effective burst size at equilibrium

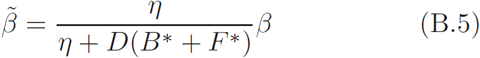

where the function 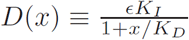 is the per capita killing rate by the maximum innate immune response when the total (uninfected and infected) bacteria density is *x*. *B*\* and *F \** are the equilibrium densities of the uninfected and infected bacteria respectively. Assuming that lysis happens at a sufficiently fast rate compared to the phage infection, the total bacteria density *B*\* + *F*\* at equilibrium can be estimated by the equilibrium bacteria density in the limit η → ∞, i.e. the equilibrium value for the model without an infected class. The value of *B*\* + *F*\* is thus approximated by the value of *B_P_* at each threshold from the original model in Eqs. (1) − (3). This results in the following scaled thresholds:

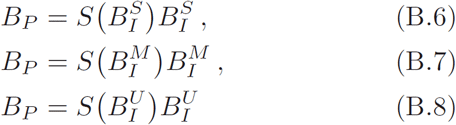

where the scaling function *S*(*B*) ≡ (1 + *D*(*B*)/*η*). The scaled thresholds are shown in Fig. B.8 and give satisfactory approximation of the transitions between regimes of phage-immune synergism, coexistence and phage extinction. The results of the analysis confirm that the same mechanism for the synergistic effect between phage and immune response applies after accounting for a reduced effective burst size.

**Figure B.8:**
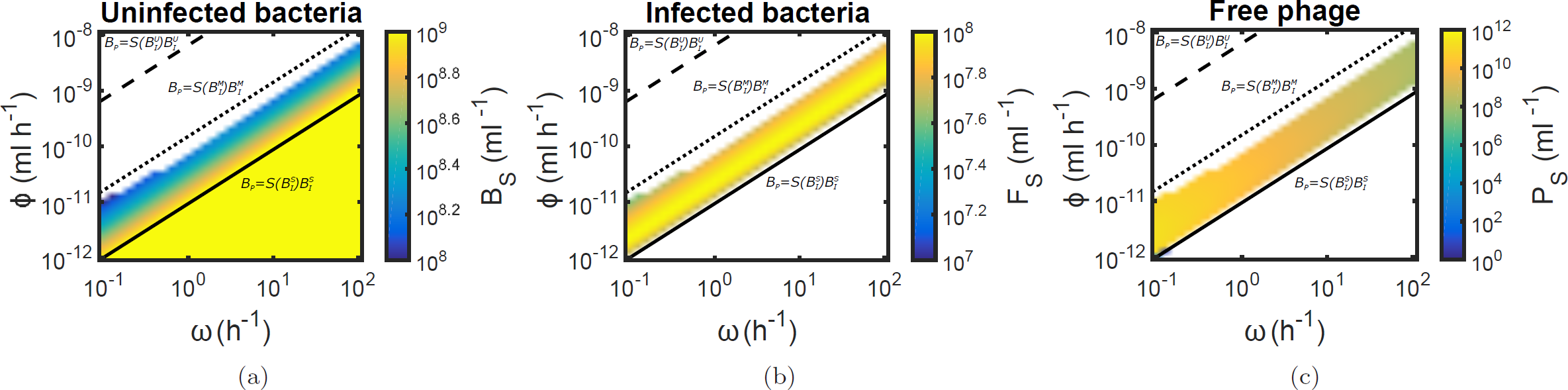
Heat map showing the dependence of (a) log steady state uninfected bacteria density *B_S_*, (b) log steady state infected bacteria density *P_S_* and (c) log steady state free phage density *P_S_* on the phage decay rate *ω* and adsorption rate *ϕ*. The white region represents the parameter space with elimination of (a) uninfected bacteria, (b) infected bacteria and (c) free phage. The solid, dotted and dashed lines are the scaled threshold relations in Eqs. (B.6), (B.7) and (B.8) respectively. The initial conditions are given by 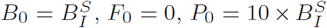 ml^−1^ and *I*_0_ = *K_I_*. The lysis rate *η* = 2 h^−1^.

## Appendix C. Effects of initial conditions on outcome of phage therapy

In this appendix we investigate the effects of initial bacterial inoculum *B*_0_ and phage dose *P*_0_ on the efficacy of phage therapy. In Fig. 6, we showed the dynamics of our model in the regimes of dynamic phage-immune synergy 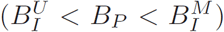, stable coexistence 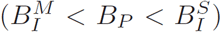 and phage extinction 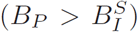. We study the robustness of these regimes to perturbations in initial conditions by modifying the initial phage dose (Fig. C.9). We find that phage-immune synergy is highly robust to variations in initial conditions, as evident by bacteria elimination even after the phage dose is reduced by five orders of magnitude (compare Fig. 6(a) with Fig. C.9(a)). In this case, the ability of phage to replicate can quickly compensate for the lower phage dose.

For phage therapy failure caused by stable coexistence between bacteria, phage and host innate immunity, our simulation shows that therapeutic efficacy may be restored by a substantial increase in phage dose (Fig. C.9(b)). This suggests that phage therapy may still be effective under limited circumstances in the stable coexistence regime, provided that a sufficiently high phage dose is used to compensate for the lack of phage-immune synergy. However, such high dosage may not be practical in clinical applications as excessive phage dosage has been linked to harmful inflammatory responses in the host (Pincus *et al*., 2015). Finally, in the phage extinction regime, even a significant increase in phage dose would not eliminate the bacteria (Fig. C.9(c)). The low phage adsorption rate in this case leads to rapid elimination of phage from the system that can negate the effect of a higher initial dose.

To verify the generality of these results, we compute the steady state bacteria density for a range of values of *B*_0_ and *P*_0_ under the three different regimes (Fig. C.10). Our results confirm the robustness of phage-immune synergy, with bacteria elimination observed over a wide range of initial conditions (Fig. C.10(a)). In the stable coexistence regime shown in Fig. C.10(b), bacteria can persist when the immune response alone is insufficient to eliminate the bacteria 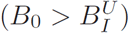. Bacteria elimination can be restored by either having a very high phage dose (> 10^10^ ml^−1^) or a high initial bacteria density (> 10^9^ ml^−1^). The paradoxical requirement of a high bacterial inoculum for subsequent bacteria elimination stems from the rapid replication of phage in the presence of high bacteria density. In such a case, phage population reaches a high level and eventually eliminate the bacteria. However, in practice the high bacterial burden may cause extensive inflammation and tissue damage, leading to host death before the phage population can reach sufficiently high densities. In the regime of phage extinction, increase in phage dose only has a marginal effect on the fate of the bacteria as the phage population cannot sustain itself (Fig. C.10(c)).

**Figure C.9:**
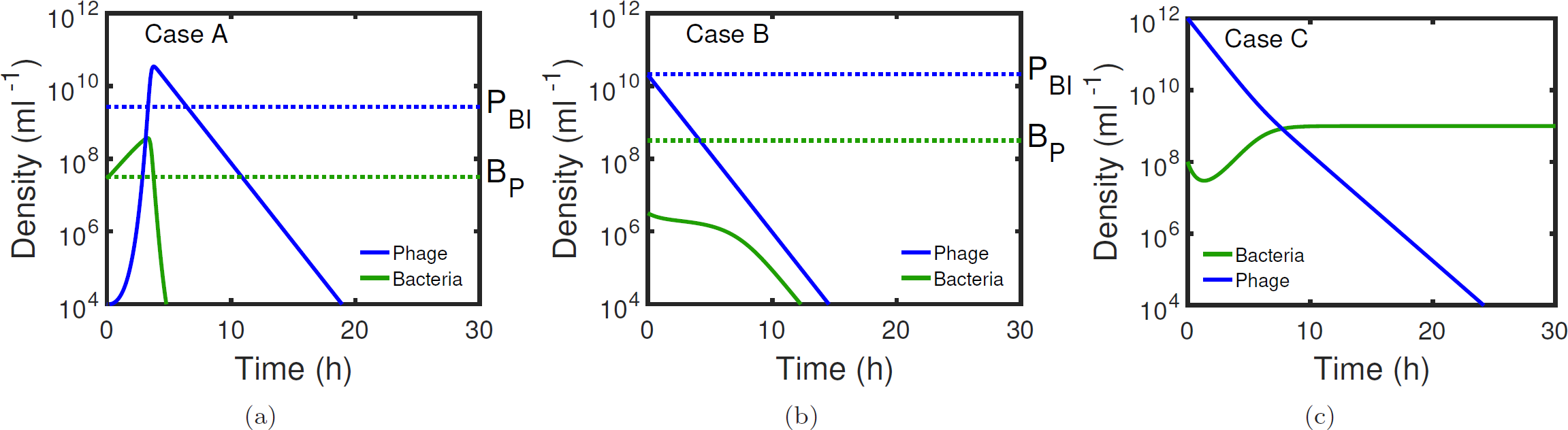
Time series of bacteria (green curve) and phage densities (blue curve) with the parameters at the three sample points A 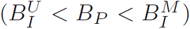, B 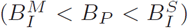 and C 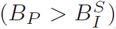 in Fig. 5. This figure is same as Fig. 6 but with different initial phage dose *P*_0_ given by (a) 10^4^ ml^−1^, (b) 2 × 10^10^ ml^−1^ and (c) 10^12^ ml^−1^.

**Figure C.10:**
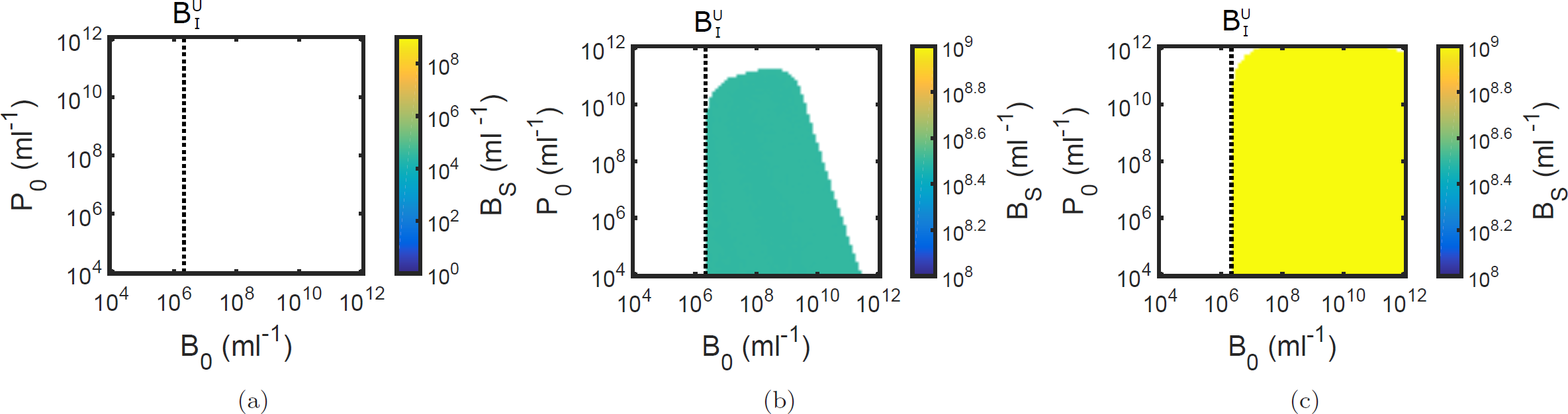
Effects of initial bacterial inoculum and phage dose on steady-state bacterial concentration *B_S_*. The parameters in (a), (b) and (c) are given by the three sample points A 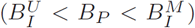, B 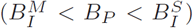 and C 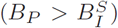 in Fig. 5 respectively. The white region represents the parameter space with elimination of bacteria. The vertical dotted line marks the value of 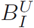.

## Acknowledgment

This work is supported by Army Research Office grant W911NF-14-1-0402. The authors thank Devika Singh for the illustration in Fig. 7. The authors thank three anonymous reviewers for feedback on the manuscript.

## Footnotes

1 https://github.com/WeitzGroup/immune-phage-synergy

## References

Abedon, S. T. (2009). Kinetics of phage-mediated biocontrol of bacteria. Foodborne Pathog Dis 6(7), 807–815.

Alemayehu, D., Casey, P. G., McAuliffe, O., Guinane, C. M., Martin, J. G., Shanahan, F., Coffey, A., Ross, R. P. & Hill, C. (2012). Bacteriophages ϕMR299–2 and ϕNH-4 can eliminate pseudomonas aeruginosa in the murine lung and on cystic fibrosis lung airway cells. mBio 3(2).

Azeredo, J. & Sutherland, I. W. (2008). The use of phages for the removal of infectious biofilms. Curr Pharm Biotechnol 9(4), 261–266.

Betts, A., Kaltz, O. & Hochberg, M. E. (2014). Contrasted co-evolutionary dynamics between a bacterial pathogen and its bacteriophages. Proc Natl Acad Sci U S A 111(30), 11109–11114.

Brockhurst, M. A. & Koskella, B. (2013). Experimental coevolution of species interactions. Trends Ecol Evol 28(6), 367–375.

Buckling, A. & Rainey, P. B. (2002). Antagonistic coevolution between a bacterium and a bacteriophage. Proc R Soc B 269(1494), 931–936.

Cairns, B. J., Timms, A. R., Jansen, V. A. A., Connerton, I. F. & Payne, R. J. H. (2009). Quantitative models of in vitro bacteriophage-host dynamics and their application to phage therapy. PLOS Pathog 5(1), 1–10.

Campbell, A. (1961). Conditions for the existence of bacteriophage. Evolution 15(2), 153–165.

Campbell, A. (2003). The future of bacteriophage biology. Nat Rev Genet 4(6), 471–477.

Capparelli, R., Nocerino, N., Iannaccone, M., Ercolini, D., Parlato, M., Chiara, M. & Iannelli, D. (2010). Bacteriophage therapy of salmonella enterica: a fresh appraisal of bacteriophage therapy. J Infect Dis 201(1), 52–61.

Carlton, R. M. (1999). Phage therapy: past history and future prospects. Arch Immunol Ther Exp 47, 267–274.

Chao, L., Levin, B. R. & Stewart, F. M. (1977). A complex community in a simple habitat: An experimental study with bacteria and phage. Ecology 58(2), 369–378.

Clark, J. R. (2015). Bacteriophage therapy: history and future prospects. Future Virol 10(4), 449–461.

Costerton, J. W., Stewart, P. S. & Greenberg, E. P. (1999). Bacterial biofilms: A common cause of persistent infections. Science 284(5418), 1318–1322.

de Kievit, T. R. & Iglewski, B. H. (2000). Bacterial quorum sensing in pathogenic relationships. Infect Immun 68(9), 4839–4849.

De Paepe, M. & Taddei, F. (2006). Viruses’ life history: towards a mechanistic basis of a trade-off between survival and reproduction among phages. PLOS Biol 4(7), e193.

Debarbieux, L., Leduc, D., Maura, D., Morello, E., Criscuolo, A., Grossi, O., Balloy, V. & Touqui, L. (2010). Bacteriophages can treat and prevent pseudomonas aeruginosa lung infections. J Infect Dis 201(7), 1096–1104.

Dennehy, J. J. (2012). What can phages tell us about host-pathogen coevolution? Int J Evol Biol 2012.

Drusano, G. L., Fregeau, C., Liu, W., Brown, D. L. & Louie, A. (2010). Impact of burden on granulocyte clearance of bacteria in a mouse thigh infection model. Antimicrob Agents Chemother 54(10), 4368–4372.

Drusano, G. L., VanScoy, B., Liu, W., Fikes, S., Brown, D. & Louie, A. (2011). Saturability of granulocyte kill of pseudomonas aeruginosa in a murine model of pneumonia. Antimicrob Agents Chemother 55(6), 2693–2695.

Ensminger, A. W., Yassin, Y., Miron, A. & Isberg, R. R. (2012). Experimental evolution of legionella pneumophila in mouse macrophages leads to strains with altered determinants of environmental survival. PLOS Pathog 8(5), 1–12.

Filippov, A. A., Sergueev, K. V., He, Y., Huang, X.-Z., Gnade, B. T., Mueller, A. J., Fernandez-Prada, C. M. & Nikolich, M. P. (2011). Bacteriophage-resistant mutants in yersinia pestis: Identification of phage receptors and attenuation for mice. PLOS ONE 6(9), 1–11.

FranÇois, B., Jafri, H. S. & Bonten, M. (2016). Alternatives to antibiotics. Intensive Care Med, 1–3.

Gellatly, S. L. & Hancock, R. E. W. (2013). Pseudomonas aeruginosa: new insights into pathogenesis and host defenses. Pathog Dis 67(3), 159–173.

Gill, J. J. & Hyman, P. (2010). Phage choice, isolation, and preparation for phage therapy. Curr Pharm Biotechnol 11(1), 2–14.

Guenther, S., Huwyler, D., Richard, S. & Loessner, M. J. (2009). Virulent bacteriophage for efficient biocontrol of listeria monocytogenes in ready-to-eat foods. Appl Environ Microbiol 75(1), 93–100.

Guo, B., Abdelraouf, K., Ledesma, K. R., Chang, K.-T., Nikolaou, M. & Tam, V. H. (2011). Quantitative impact of neutrophils on bacterial clearance in a murine pneumonia model. Antimicrob Agents Chemother 55(10), 4601–4605.

GutiÉrrez, D., Vandenheuvel, D., MartÍnez, B., RodrÍguez, A., Lavigne, R. & GarcÍa, P. (2015). Two phages, phiIPLA-RODI and phiIPLA-C1C, lyse monoand dual-species staphylococcal biofilms. Appl Environ Microbiol 81(10), 3336–3348.

Hagens, S. & Loessner, M. J. (2007). Application of bacteriophages for detection and control of foodborne pathogens. Appl Microbiol Biotechnol 76(3), 513–519.

Hall, A. R., Scanlan, P. D., Morgan, A. D. & Buckling, A. (2011). Host-parasite coevolutionary arms races give way to fluctuating selection. Ecol Lett 14(7), 635–642.

Hermoso, J. A., GarcÍa, J. L. & GarcÍa, P. (2007). Taking aim on bacterial pathogens: from phage therapy to enzybiotics. Curr Opin Microbiol 10(5), 461–472.

Hodyra-Stefaniak, K., Miernikiewicz, P., DrapaŁ, J., Drab, M., JoŃczyk-Matysiak, E., Lecion, D., KaŹmierczak, Z., Beta, W., Majewska, J., Harhala, M. et al. (2015). Mammalian host-versus-phage immune response determines phage fate in vivo. Sci Rep 5, 14802.

Hyman, P. & Abedon, S. T. (2010). Bacteriophage host range and bacterial resistance. In: Advances in Applied Microbiology, vol. 70 of Advances in Applied Microbiology. Academic Press, pp. 217–248.

Jubrail, J., Morris, P., Bewley, M. A., Stoneham, S., Johnston, S. A., Foster, S. J., Peden, A. A., Read, R. C., Marriott, H. M. & Dockrell, D. H. (2016). Inability to sustain intraphagolysosomal killing of staphylococcus aureus predisposes to bacterial persistence in macrophages. Cell Microbiol 18(1), 80–96.

Khan Mirzaei, M. & Nilsson, A. S. (2015). Isolation of phages for phage therapy: A comparison of spot tests and efficiency of plating analyses for determination of host range and efficacy. PLOS ONE 10(3), 1–13.

Kingwell, K. (2015). Bacteriophage therapies re-enter clinical trials. Nat Rev Drug Discov 14(8), 515–516.

Labrie, S. J., Samson, J. E. & Moineau, S. (2010). Bacteriophage resistance mechanisms. Nature Rev Microbiol 8(5), 317–327.

Leggett, H. C., Cornwallis, C. K. & West, S. A. (2012). Mechanisms of pathogenesis, infective dose and virulence in human parasites. PLOS Pathog 8(2), 1–5.

Lenski, R. E. (1984). Coevolution of bacteria and phage: Are there endless cycles of bacterial defenses and phage counterdefenses? J Theor Biol 108(3), 319–325.

Leverentz, B., Conway, W. S., Camp, M. J., Janisiewicz, W. J., Abuladze, T., Yang, M., Saftner, R. & Sulakvelidze, A. (2003). Biocontrol of listeria monocytogenes on fresh-cut produce by treatment with lytic bacteriophages and a bacteriocin. Appl Environ Microbiol 69(8), 4519–4526.

Levin, B. R. & Bull, J. J. (1996). Phage therapy revisited: The population biology of a bacterial infection and its treatment with bacteriophage and antibiotics. Am Nat 147(6), 881–898.

Levin, B. R. & Bull, J. J. (2004). Population and evolutionary dynamics of phage therapy. Nature Rev Microbiol 2(2), 166–173.

Levin, B. R., Stewart, F. M. & Chao, L. (1977). Resource-limited growth, competition, and predation: A model and experimental studies with bacteria and bacteriophage. Am Nat 111(977), 3–24.

Li, Y., Karlin, A., Loike, J. D. & Silverstein, S. C. (2002). A critical concentration of neutrophils is required for effective bacterial killing in suspension. Proc Natl Acad Sci U S A 99(12), 8289–8294.

Loc-Carrillo, C. & Abedon, S. T. (2011). Pros and cons of phage therapy. Bacteriophage 1(2), 111–114.

Meyer, J. R., Dobias, D. T., Weitz, J. S., Barrick, J. E., Quick, R. T. & Lenski, R. E. (2012). Repeatability and contingency in the evolution of a key innovation in phage lambda. Science 335(6067), 428–432.

Nau, G. J., Richmond, J. F. L., Schlesinger, A., Jennings, E. G., Lander, E. S. & Young, R. A. (2002). Human macrophage activation programs induced by bacterial pathogens. Proc Natl Acad Sci U S A 99(3), 1503–1508.

Payne, R. J. H. & Jansen, V. A. A. (2000). Phage therapy: The peculiar kinetics of self-replicating pharmaceuticals. Clin Pharmacol Ther 68(3), 225–230.

Payne, R. J. H. & Jansen, V. A. A. (2001). Understanding bacteriophage therapy as a density-dependent kinetic process. J Theor Biol 208(1), 37–48.

Payne, R. J. H. & Jansen, V. A. A. (2003). Pharmacokinetic principles of bacteriophage therapy. Clinical Pharmacokinetics 42(4), 315–325.

Pincus, N. B., Reckhow, J. D., Saleem, D., Jammeh, M. L., Datta, S. K. & Myles, I. A. (2015). Strain specific phage treatment for staphylococcus aureus infection is influenced by host immunity and site of infection. PLoS ONE 10(4), 1–16.

Potera, C. (2013). Phage renaissance: new hope against antibiotic resistance. Environ Health Persp 121(2), a48.

Reardon, S. (2014). Phage therapy gets revitalized. Nature 510(7503), 15–16.

Reutershan, J., Basit, A., Galkina, E. V. & Ley, K. (2005). Sequential recruitment of neutrophils into lung and bronchoalveolar lavage fluid in lps-induced acute lung injury. Am J Physiol Lung Cell Mol Physiol 289(5), L807–L815.

Rodney, M. D. (2009). Preventing biofilms of clinically relevant organisms using bacteriophage. Trends Microbiol 17(2), 66–72.

Sadek, M. I., Sada, E., Toossi, Z., Schwander, S. K. & Rich, E. A. (1998). Chemokines induced by infection of mononuclear phagocytes with mycobacteria and present in lung alveoli during active pulmonary tuberculosis. Am J Respir Cell Mol Biol 19(3), 513–521.

Seed, K. D., Yen, M., Shapiro, B. J., Hilaire, I. J., Charles, R. C., Teng, J. E., Ivers, L. C., Boncy, J., Harris, J. B. & Camilli, A. (2014). Evolutionary consequences of intra-patient phage predation on microbial populations. eLife 3, e03497.

Semler, D. D., Goudie, A. D., Finlay, W. H. & Dennis, J. J. (2014). Aerosol phage therapy efficacy in burkholderia cepacia complex respiratory infections. Antimicrob Agents Chemother 58(7), 4005–4013.

Seo, J., Seo, D. J., Oh, H., Jeon, S. B., Oh, M.-H. & Choi, C. (2016). Inhibiting the growth of escherichia coli o157:h7 in beef, pork, and chicken meat using a bacteriophage. Korean J Food Sci An 2(2).

Shao, Y. & Wang, I.-N. (2008). Bacteriophage adsorption rate and optimal lysis time. Genetics 180(1), 471–482.

Skurnik, M., Pajunen, M. & Kiljunen, S. (2007). Biotechnological challenges of phage therapy. Biotechnol Lett 29(7), 995–1003.

Smith, A. M. & Smith, A. P. (2016). A critical, nonlinear threshold dictates bacterial invasion and initial kinetics during influenza. Sci Rep 6, 38703.

Strogatz, S. H. (2014). Nonlinear dynamics and chaos: with applications to physics, biology, chemistry, and engineering. Boulder, Colorado: Westview Press.

Sully, E. K., Malachowa, N., Elmore, B. O., Alexander, S. M., Femling, J. K., Gray, B. M., DeLeo, F. R., Otto, M., Cheung, A. L., Edwards, B. S., Sklar, L. A., Horswill, A. R., Hall, P. R. & Gresham, H. D. (2014). Selective chemical inhibition of agr quorum sensing in staphylococcus aureus promotes host defense with minimal impact on resistance. PLOS Pathog 10(6), e1004174.

Summers, C., Rankin, S. M., Condliffe, A. M., Singh, N., Peters, A. M. & Chilvers, E. R. (2010). Neutrophil kinetics in health and disease. Trends Immunol 31(8), 318–324.

Thiel, K. (2004). Old dogma, new tricks–21st century phage therapy. Nature Biotechnol 22(1), 31–36.

Timothy, K. L. & Michael, S. K. (2011). The next generation of bacteriophage therapy. Curr Opin Microbiol 14(5), 524–531.

Tiwari, B. R., Kim, S., Rahman, M. & Kim, J. (2011). Antibacterial efficacy of lytic pseudomonas bacteriophage in normal and neutropenic mice models. J Microbiol 49(6), 994–999.

Trigo, G., Martins, T. G., Fraga, A. G., Longatto-Filho, A., Castro, A. G., Azeredo, J. & Pedrosa, J. (2013). Phage therapy is effective against infection by mycobacterium ulcerans in a murine footpad model. PLoS Negl Trop Dis 7(4), 1–10.

Weitz, J. S. (2015). Quantitative Viral Ecology: Dynamics of Viruses and Their Microbial Hosts. Princeton, New Jersey: Princeton University Press.

Weitz, J. S., Hartman, H. & Levin, S. A. (2005). Coevolutionary arms races between bacteria and bacteriophage. Proc Natl Acad Sci U S A 102(27), 9535–9540.

Wigginton, J. E. & Kirschner, D. (2001). A model to predict cellmediated immune regulatory mechanisms during human infection with mycobacterium tuberculosis. J Immunol 166(3), 1951–1967.

